# *Frem1* activity regulated by Sonic Hedgehog signaling in the cranial neural crest mesenchyme guides midfacial morphogenesis

**DOI:** 10.1101/2022.07.16.500318

**Authors:** Matthew T. McLaughlin, Miranda R. Sun, Tyler G. Beames, Austin C. Steward, Joshua W. M. Theisen, Hannah M. Chung, Joshua L. Everson, Ivan P. Moskowitz, Michael D. Sheets, Robert J. Lipinski

**Affiliations:** Department of Comparative Biosciences, School of Veterinary Medicine, University of Wisconsin-Madison, Madison, WI, 53706, United States of America; Department of Pediatrics, Pathology, Human Genetics and Genetic Medicine, The University of Chicago, Chicago, IL 60637; Department of Biomolecular Chemistry, School of Medicine and Public Health, University of Wisconsin-Madison, Madison, WI

**Keywords:** Frem1, Sonic hedgehog, cranial neural crest, facial morphogenesis, midface hypoplasia

## Abstract

The Frem/Fras family of extracellular matrix proteins has been linked to human face shape variation and malformation, but little is known about their regulation and biological roles in facial development. During midfacial morphogenesis in mice, we observed *Frem1* expression in the embryonic growth centers that form the median upper lip, nose, and palate. Expansive spatial gradients of *Frem1* expression in the cranial neural crest cell (cNCC) mesenchyme of these tissues suggested transcriptional regulation by a secreted morphogen. Accordingly, *Frem1* expression paralleled that of the conserved Sonic Hedgehog (Shh) target gene *Gli1* in the cNCC mesenchyme. Suggesting direct transcriptional regulation by Shh signaling, we found that *Frem1* expression is induced by SHH ligand stimulation or downstream pathway activation in cNCCs and observed GLI transcription factor binding at the *Frem1* transcriptional start site during midfacial morphogenesis. Shh pathway antagonism reduced *Frem1* expression during pathogenesis of midfacial hypoplasia, and FREM1 was sufficient to induce cNCC proliferation in a concentration-dependent manner. These findings provide novel insight into the mechanism by which the Shh pathway drives midfacial morphogenesis and reveal a functional role for *Frem1* in cNCC biology that establishes the developmental basis for *FREM1-*associated face shape variation and malformation.

## Introduction

Substantial variation in facial morphology exists within and across human populations, and approximately one-third of all human birth defects involve craniofacial anomalies [8; 34; 53]. The Frem/Fras family of extracellular matrix proteins has been implicated in face shape variation and overt facial malformations. Mutations in human *FREM1, FREM2*, and *FRAS1* genes cause Manitoba-oculo-tricho-anal (MOTA), bifid nose with or without anorectal and renal anomalies (BNAR), and Fraser syndromes [3; 35; 44]. Craniofacial anomalies described in these syndromes include bifid nose, thin upper lip, shortened philtrum, and orofacial clefting [3; 7; 13; 45]. Additionally, genome-wide association studies of human facial morphology have linked *FREM1* polymorphisms to shape variation of the central upper lip, while varying degrees of midfacial asymmetry and hypoplasia have been described in *Frem1* mutant mice [4; 26; 52; 53]. Despite these genotype-phenotype relationships from human and mouse studies, the regulatory mechanisms and roles of *Frem1* in craniofacial morphogenesis have not been reported.

The biological activity of the Frem/Fras family has been most extensively characterized in epidermal development. FREM1 secreted by dermis mesenchymal cells is thought to form a ternary complex in the basement membrane with epidermally-secreted FREM2 and FRAS1 [21; 39]. The role of this complex in cross-linking the epidermal basement membrane to the developing dermis manifests as a characteristic “blebbing” phenotype described in *Frem/Fras* mutant mice and related human syndromes [21; 47]. However, an exclusive role in epidermal development is unlikely to explain the facial dysmorphology and malformations linked to Frem/Fras genetic variation.

In this study, we generated a detailed spatiotemporal expression profile of *Frem1* during mouse embryonic facial morphogenesis, defined upstream *Frem1* regulatory mechanisms, and examined downstream influences of FREM1 on cranial neural crest cell (cNCC) biology. We found that transcriptional regulation by Sonic Hedgehog (Shh) signaling establishes spatial gradients of *Frem1* in the cNCC mesenchyme, that FREM1 promotes concentration-dependent cNCC proliferation, and that Shh-Frem1 signaling is disrupted during pathogenesis of midface hypoplasia. These observations establish previously unrecognized regulators and roles of *Frem1* in cNCC biology that provide new insight into the mechanisms underlying variation in midfacial morphogenesis.

## Results

Midfacial morphogenesis involves orchestrated outgrowth and fusion of paired facial growth centers (Fig. 1a-b) that are comprised of cNCC-derived mesenchyme covered by surface ectoderm (Fig. 1c-f). We applied microdissection, enzymatic separation, and qPCR to assess tissue-specific gene expression at gestational day (GD)11, when the medial nasal processes (MNPs) and the maxillary processes (MxPs) fuse bilaterally to close the upper lip in the mouse. In both of these tissues, *Frem2* and *Fras1* were predominantly detected in the epithelium, while *Frem1* expression was enriched in the mesenchyme (Fig. 1g-h). Whole-mount *in situ* hybridization revealed *Frem1* expression in the facial growth centers, with strong staining apparent in both the MNPs and MxPs (Fig. 1i). This was confirmed by staining of tissue sections, which revealed expansive spatial gradients of *Frem1* expression in MNP and MxP mesenchyme (Fig. 1j). *Frem2* expression was observed in the ectodermal epithelium surrounding the growth centers and lining the entire nasal pit, while *Fras1* expression appeared limited to the epithelium on the medial aspect of the nasal pit (Fig. 1k-l).

**Figure 1.**
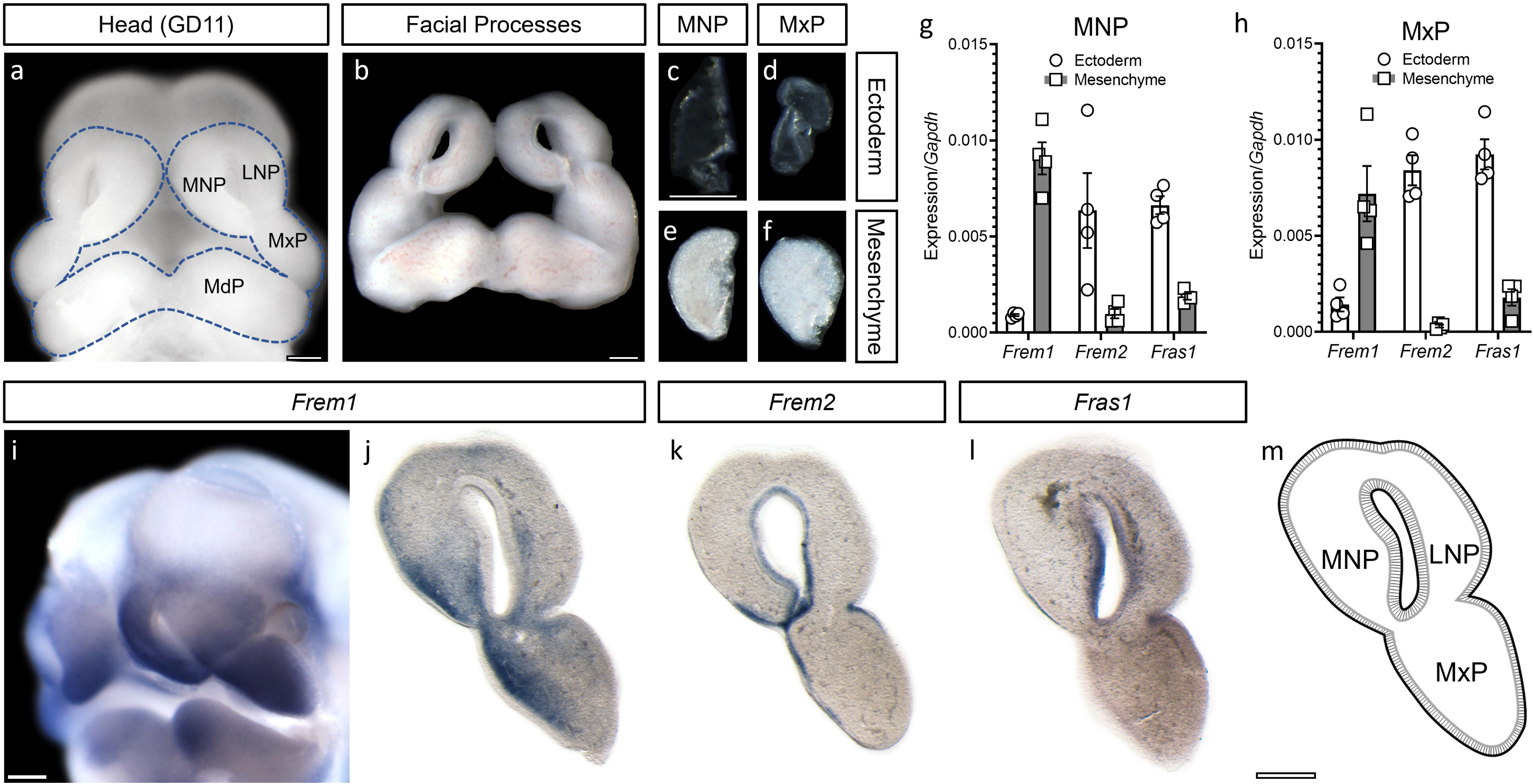
*Frem1* is expressed in the cNCC mesenchyme during midfacial morphogenesis. The bilaterally paired facial growth centers that form the midface are shown in an intact GD11.0 embryo (a), after microdissection (b), and after enzymatic digestion to separate the ectoderm and cNCC-derived mesenchyme (c-f). Gene expression in ectodermal and mesenchymal compartments of GD11 MNP and MxP tissue was determined by qPCR (g-h). Individual values are plotted along with the mean ± SEM of four independently collected and pooled tissue samples. Whole-mount tissue (i) and sections through the facial growth centers (j-l) were stained by ISH to visualize expression of *Frem1* (i-j), *Frem2* (k), and *Fras1* (l). At GD11, the facial growth centers fuse to form a lambdoidal junction as shown in schematic (m). MNP, medial nasal process; MxP, maxillary processes; LNP, lateral nasal process. Scale bars: 0.25 mm.

The observed spatial gradient of *Frem1* expression prompted us to examine potential regulation by Sonic Hedgehog (SHH), a morphogen and critical regulator of facial development [10; 15; 24]. SHH ligand secreted by the facial ectodermal epithelium establishes a morphogen gradient that induces pathway activity in the cNCC mesenchyme [18], which can be visualized as expression of the conserved pathway target gene, *Gli1*. We therefore examined expression of *Frem1* and *Gli1* at key stages of midfacial morphogenesis. *Frem1* and *Gli1* were expressed in overlapping spatial gradients in the cNCC-derived mesenchyme of the MNPs throughout their expansion from GD10.25 to GD11 (Fig. 2a-c, g-i). Overlapping gradients of *Frem1* and *Gli1* were also observed in the mesenchyme of the MxPs at GD11, in MxP-derived palatal shelf mesenchyme, and in the mandibular processes (MdP) at GD13 (Fig. 2d-f, j-l). In each of these domains, *Frem1* expression appeared restricted to the mesenchyme and concomitant with *Gli1*, while *Gli1* staining was also detected in regions of neuroectoderm (Fig. 2a,d).

**Figure 2.**
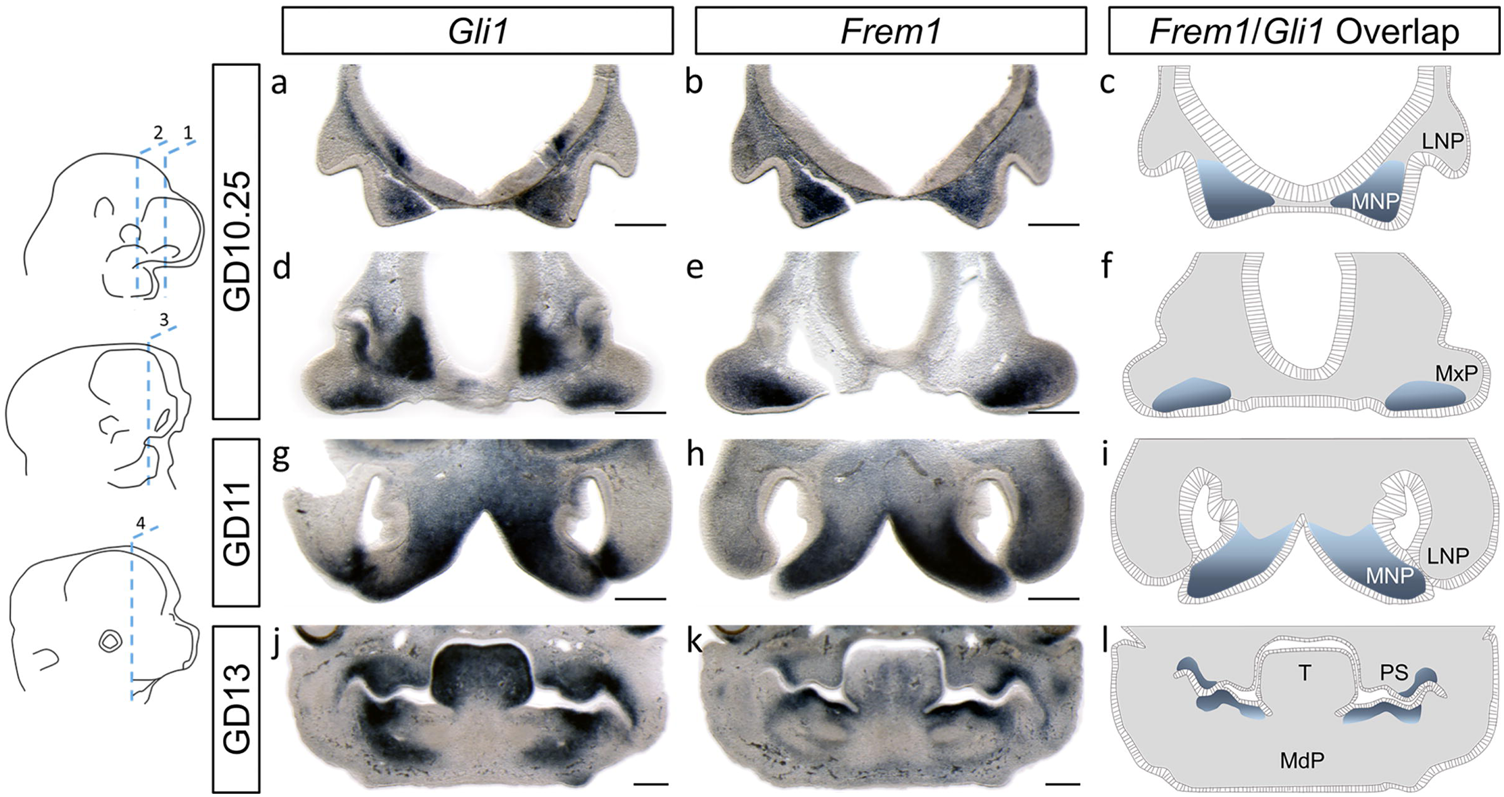
*Frem1* expression overlaps with Shh pathway target *Gli1* during midfacial morphogenesis. The spatial expression of *Frem1* and *Gli1* by was assessed by ISH at key stages of midfacial morphogenesis. Embryonic stages and planes of section are depicted by schematics in the left column. GD10.25 embryos were sectioned to visualize nascent MNP, LNP (a-c, plane of section 1) or MxP tissues (d-f, plane of section 2). GD11 embryos were sectioned to visualize MNP and LNP tissues along their proximal to distal axes (g-i, plane of section 3). GD13 embryos were sectioned to visualize the MxP-derived palatal shelves situated vertically along the sides of the tongue (T) (j-l, plane of section 4). Staining is shown on adjacent sections for each stage/plane of section. Areas of apparent overlap in *Gli1* and *Frem1* expression are shown in schematics (c,f,i,l). MNP, medial nasal process; MxP, maxillary processes; LNP, lateral nasal process; MdP, mandibular process; MdP, mandibular process. Scale bar: 0.25 mm.

We next examined whether Shh signaling directly regulates *Frem1* expression in a mouse cNCC line (O9-1) that recapitulates the expression signature and differentiation capacity of *in vivo* multipotent cNCCs [17]. Stimulation of cultured cNCCs with SHH ligand significantly increased the expression of both *Gli1* and *Frem1* compared to vehicle alone (Fig. 3a-b). The SHH ligand-induced expression of both *Gli1* and *Frem1* was blocked by exposure to vismodegib, a specific inhibitor of the obligate Shh pathway signal transducing protein Smoothened (SMO) [41]. We therefore tested whether genetic and pharmacologic activation of SMO could directly regulate *Frem1* expression. Overexpression of a constitutively active form of human *SMO* in cultured cNCCs resulted in significantly increased *Gli1* and *Frem1* expression (Fig. 3c-d). Accordingly, acute pathway activation via addition of the small molecule SMO activator SAG was sufficient to upregulate *Gli1* and *Frem1* expression (Fig. 3e-f).

**Figure 3.**
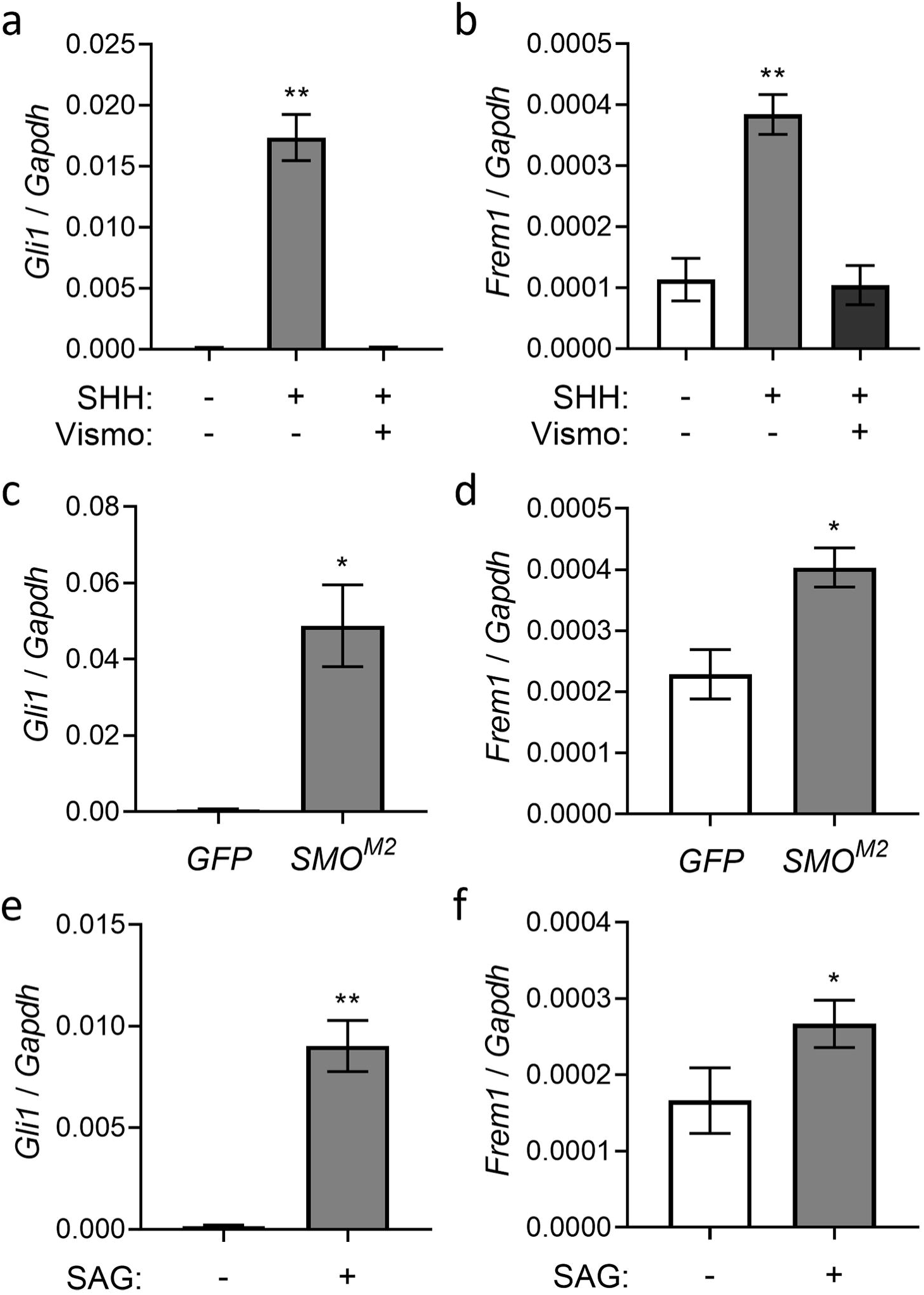
*Frem1* is regulated by the Shh pathway in cNCCs. cNCCs were cultured with or without SHH ligand (0.4 µg/mL) and with or without the Smoothened antagonist vismodegib (Vismo, 100 nM) (a-b). SHH ligand caused an increase in *Gli1* and *Frem1* expression, which was blocked by the addition of vismodegib. Expression of *Gli1* and *Frem1* are increased in cNCCs expressing a constitutively active form of human Smoothened (*SMO*^*M2*^) relative to a GFP expressing line (c-d). cNCCs cultured with the Smoothened agonist SAG (50 nM) similarly demonstrated increased expression of *Gli1* and *Frem1* compared to vehicle alone (e-f). Values represent the mean ± SEM of N=5 biological replicates for each condition. *p<0.05, ** p<0.01 (ANOVA with Tukey’s post hoc test or two-tailed t-test).

Our observations from *in vivo* development and cNCC culture suggested transcriptional regulation of *Frem1* by the Shh signaling pathway. Three zinc finger GLI proteins, GLI1, GLI2, and GLI3, regulate transcription of Shh target genes by binding to consensus GLI DNA sequence motifs [20; 55]. To identify GLI binding sites in the developing face, we utilized published mouse GLI3 ChIP-seq data sets generated from GD11.5 whole face (including MNP and MxP) and isolated MdP tissue [9]. In addition to *Frem1* and *Gli1*, we examined *Ccnd2*, an established transcriptional target of Shh-Gli [25; 36; 55]. ChIP-seq revealed binding of GLI3 at the transcription start sites of *Frem1, Gli1*, and *Ccnd2* (Fig. 4), suggesting GLI transcription factors directly regulate *Frem1* expression during facial morphogenesis.

**Figure 4.**
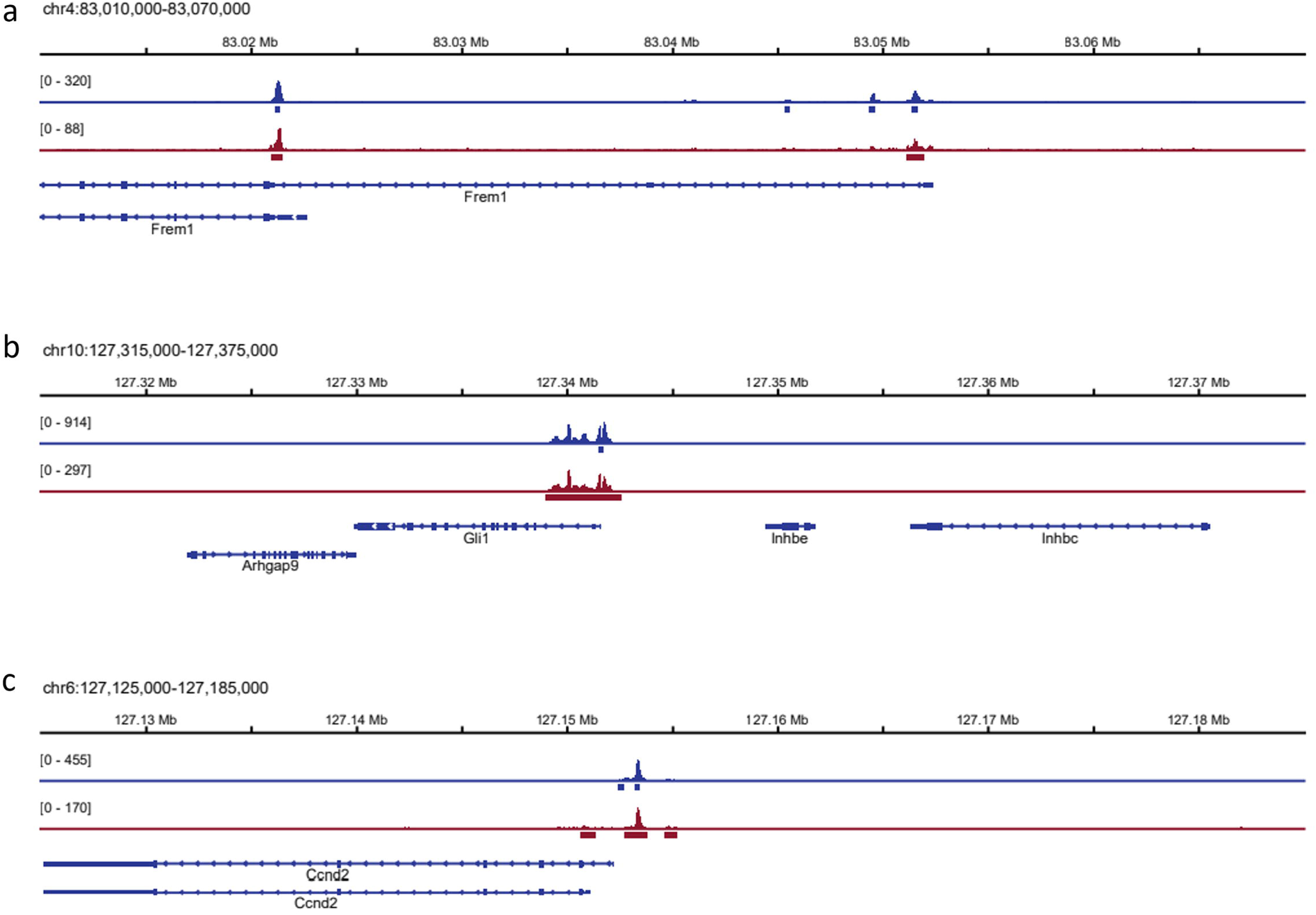
GLI binding to the *Frem1* transcriptional start site during facial morphogenesis. GLI3 ChIP-seq data from GD11.5 whole face (MNP, MxP, LNP, MdP, blue tracks) and isolated MdP tissue (red tracks) were analyzed for GLI3 binding at Shh pathway target genes. GLI3 peaks (blue or red rectangles below ChIP-seq tracks) were present at both RefSeq [38] annotated transcription start sites for *Frem1* (a). GLI3 ChIP-seq signal at the *Frem1* promoters was comparable to GLI3 signal at the promoters of known Shh targets *Gli1* and *Ccnd2* (b-c).

*Frem1* mutant mice exhibit varying degrees of midfacial hypoplasia and asymmetry [52]. We have previously demonstrated that *in utero* antagonism of the Shh signaling pathway disrupts midfacial morphogenesis, resulting in a spectrum of related phenotypic outcomes ranging from midfacial hypoplasia to cleft lip (Fig. 5a-c) [29; 30]. We therefore examined the impact of Shh pathway antagonism on *Frem1* expression during the pathogenesis of midfacial hypoplasia/cleft lip. Embryos were exposed *in utero*, beginning at GD8.25, to the Shh pathway inhibitor cyclopamine or vehicle alone and collected at GD9.25 during initial pathogenesis [10]. Frontonasal prominence (FNP) tissue that gives rise to the paired medial and lateral nasal processes was isolated by microdissection, and gene expression was assessed by qPCR. Expression levels of both *Gli1* and *Frem1* were significantly reduced in FNP tissue of embryos exposed to cyclopamine compared to vehicle alone (Fig. 5d). Spatial gene expression was then examined on a parallel cohort of embryos collected at GD10, when the MNPs are undergoing initial outgrowth. Consistent with observations shown in Fig. 2, both *Frem1* and *Gli1* were expressed in spatial gradients in the MNP mesenchyme of vehicle-exposed embryos (Fig. 5e, g). In embryos exposed to the Shh pathway antagonist cyclopamine, mesenchymal expression of both *Gli1* and *Frem1* was markedly diminished, while *Gli1* staining in the neuroectoderm was retained (Fig. 5 f,h).

**Figure 5.**
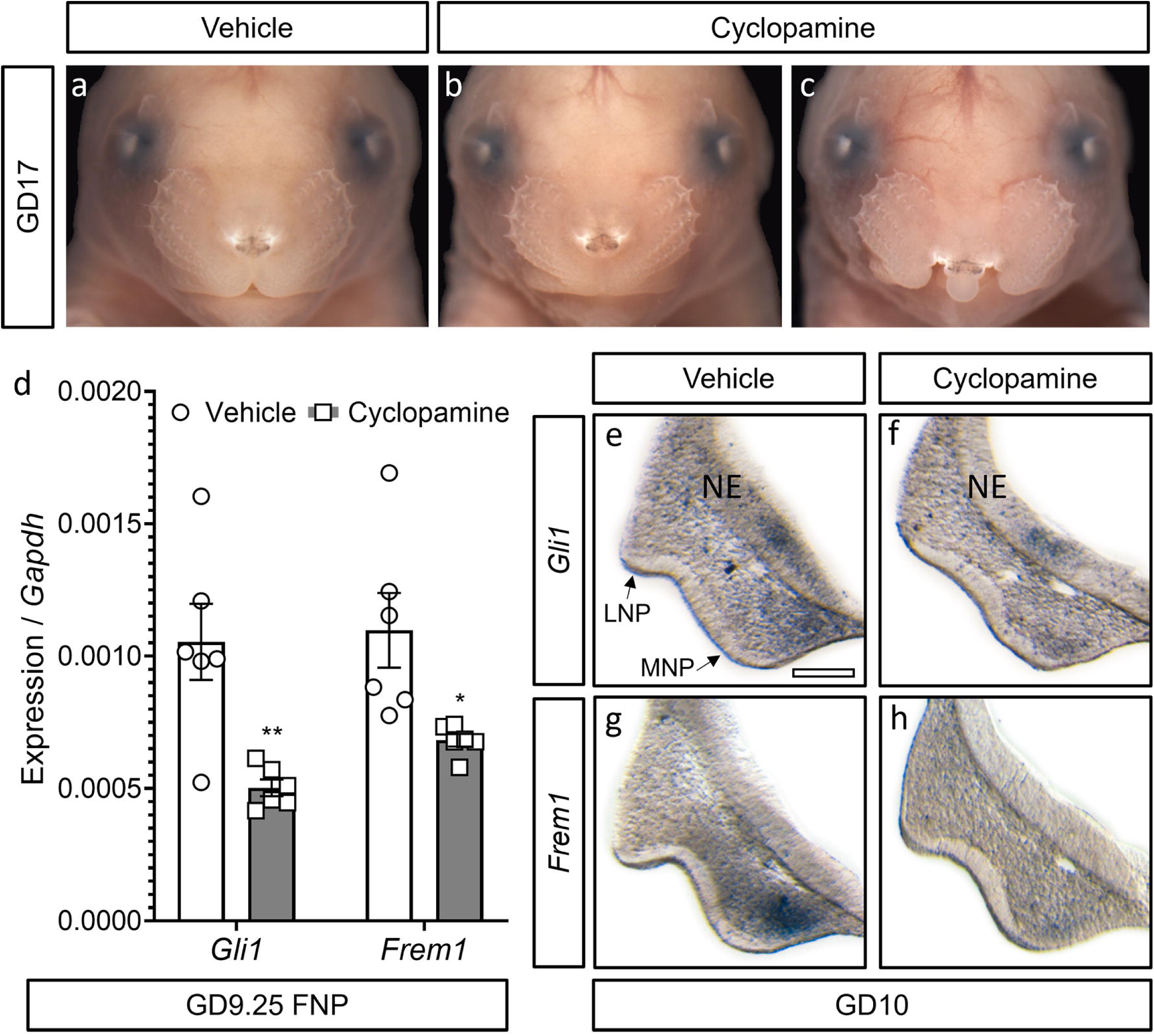
*Frem1* expression is diminished by Shh pathway inhibition during pathogenesis of midfacial hypoplasia/cleft lip. *In utero* exposure to the Shh pathway antagonist cyclopamine from GD8.25 to 9.5 disrupts midfacial morphogenesis, resulting in either midfacial hypoplasia (b) or cleft lip (c) compared to control (a) as shown at GD17. Frontonasal prominence (FNP) tissue that gives rise to the MNPs and LNPs was microdissected from vehicle- and cyclopamine-exposed embryos at GD9.25, and gene expression was determined by qPCR (d). Expression data from pooled tissue of an entire litter are shown as individual points along with mean ± SEM. Expression of both *Gli1* and *Frem1* was significantly reduced in FNP tissue from embryos exposed to cyclopamine versus vehicle alone. * p<0.05, ** p<0.01 (two-tailed t-test). Sections through MNP and LNP tissue produced from GD10 vehicle- and cyclopamine-exposed embryos were stained by ISH to visualize expression of *Gli1* (e-f) and *Frem1* (g-h). Cyclopamine exposure reduced expression of *Gli1* and *Frem1* in the MNP mesenchyme. FNP, frontonasal prominence; MNP, medial nasal process; LNP, lateral nasal process; NE, neuroectoderm. Scale bar: 0.125 mm.

We next assessed the influence of FREM1 on cNCC migration and proliferation. After undergoing epithelial-to-mesenchymal transition, cNCCs migrate away from the dorsal margins of the neural folds into the field of facial morphogenesis, then rapidly proliferate to form the mesenchyme of the facial growth centers. To assess migration, scratch assays were conducted with cNCCs cultured in the absence or presence of recombinant human FREM1 protein, and the widths of scratches were measured every hour for eight hours (Fig. S1). No significant impact of FREM1 was observed on the rate of cNCC migration (Fig. 6a-g). The potential influence on proliferation was then examined by culturing cNCCs in the absence or presence of graded concentrations of FREM1 for 24 hours, with EdU added to culture media for the last 2 hours. Addition of FREM1 resulted in a concentration-dependent increase in EdU incorporation (Fig. 6h-j), with the highest tested FREM1 concentration of 7.5 µg/ml resulting in a 20% increase. hese observations indicate that Frem1 potentiates cNCC proliferation, but not migration, providing a mechanism for the requirement for *Frem1* in craniofacial morphogenesis.

**Figure 6.**
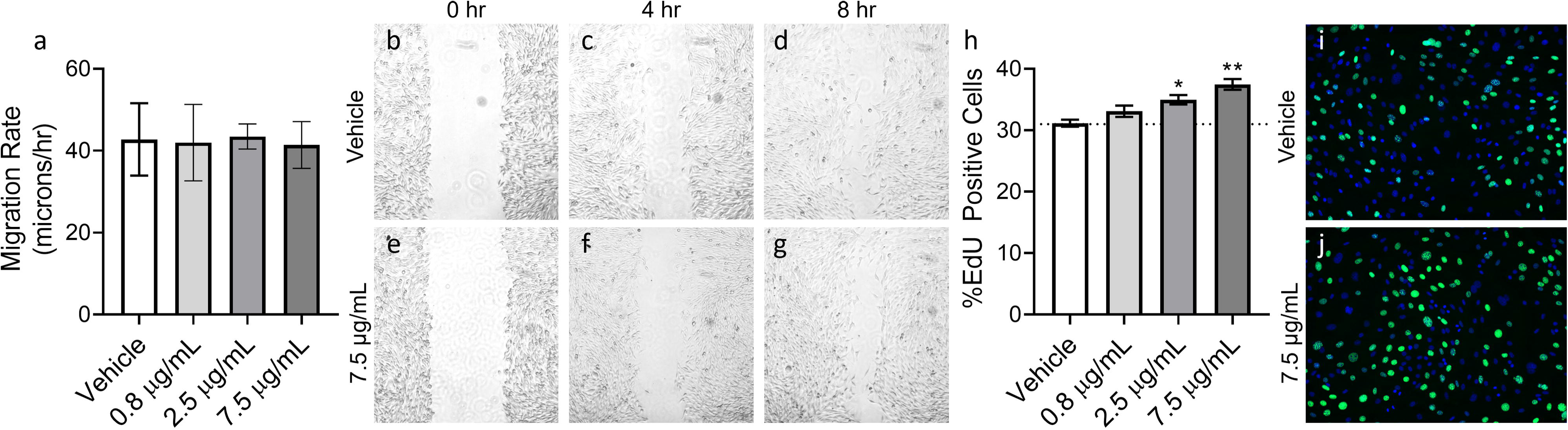
FREM1 promotes concentration-dependent cNCC proliferation. Scratch assays were performed on O9-1 cells ± recombinant FREM1, and migration rate was determined over the period of scratch closure for up to 8 hours. FREM1 had no impact on the rate of O9-1 cell migration at any concentration tested (a). Representative images from vehicle- and 7.5 μg/mL FREM1-treated cells show scratch closure over time (b-g). EdU incorporation was used to assess cell proliferation in FREM1-treated O9-1 cells. Addition of recombinant FREM1 protein to culture media increased proliferation in a concentration-dependent manner (h). Representative images of vehicle- and FREM1-treated O9-1 cells stained with Hoechst (blue) and EdU (green) (i-j). Values represent the mean ± SEM of N=3 biological replicates for each condition. *p<0.05, **p<0.01 (one-way ANOVA).

## Discussion

This study is the first to demonstrate that *Frem1* is regulated by the Shh signaling pathway, a key driver of facial morphogenesis [18; 33]. Previous studies have described *Frem1* expression in restricted domains localized to sites of epithelial-mesenchymal interaction in several developmental contexts [3; 46]. In the embryonic facial growth centers, we observed expansive *Frem1* spatial gradients that parallel *Gli1*, a reliable indicator of Shh pathway activity. The canonical Shh signaling pathway is initiated by SHH ligand binding to the transmembrane protein PTCH1, relieving its repression of SMO, which then localizes to the primary cilium and triggers a downstream signaling cascade culminating in transcriptional regulation of pathway target genes by the GLI transcription factor family. We found that SHH ligand-stimulated upregulation of *Frem1* can be blocked by SMO inhibition and that genetic or pharmacologic SMO activation is sufficient to induce *Frem1* expression. Leveraging a previously published GLI3 ChIP-seq data set [9], we also found evidence of GLI transcription factor binding at the *Frem1* promoter *in vivo*. Taken together, these findings suggest that *Frem1* expression is directly regulated by canonical Shh-Gli signaling during midfacial morphogenesis.

The Shh-Frem1 regulatory relationship revealed in this study contextualizes *Frem1*-associated facial phenotypes with those resulting from Shh pathway inhibition. Mutations in *FREM1* result in BNAR syndrome (OMIM #608980) and MOTA syndrome (OMIM #248450). Facial features described in affected individuals include bifid or bulbous nasal tip, short philtrum, thin upper lip, highly arched palate, and upper incisor abnormalities [2; 3; 6; 12; 44]. *FREM1* polymorphisms have also been linked to shape variation of the central upper lip [26; 53]. These outcomes are consistent with the prominent gradients of *Frem1* observed during expansion of the MNPs, which form the median aspect of the nose, including the nasal septum and nasal tip, the upper lip philtrum, and the central portion of the alveolar ridge that contains the upper central incisors. Expansion of the MNPs is dependent upon Shh signaling and highly sensitive to pathway disruption [31]. Human mutations in *SHH* and other genes encoding pathway effectors are associated with holoprosencephaly, a malformation of the developing forebrain. Reflecting the requirement of Shh activity in the development of the forebrain and adjacently developing face, the prosencephalic anomalies that define holoprosencephaly co-occur with midfacial deficiency manifesting as severe hypoplasia, median and lateral orofacial clefts, highly arched palate, and a single central incisor [43; 48; 49]. These comparisons suggest that disruption of *Frem1* itself results in outcomes that fall within, but are generally less severe than, those caused by complete Shh pathway inhibition. This premise is further supported by our observation that *Frem1* expression is reduced following Shh pathway inhibition and during pathogenesis of midfacial hypoplasia/cleft lip. *Frem1* then appears to be one of several Shh pathway target genes, including previously identified members of the Forkhead box transcription factor family, that influence cNCC biology and are individually required for midfacial morphogenesis [10; 18].

Whether *Frem1* regulation by the Shh pathway as demonstrated in cNCCs during facial development is operational in other developmental contexts is unclear. Known roles of Shh signaling do not readily correspond to the diaphragmatic and kidney malformations described to result from *Frem1* disruption [5; 19]. However, evidence suggests a potential interaction between Shh signaling and *Frem1* activity in skin and anorectal development. While the *Frem1* expression pattern in the latter developmental context has not been documented, genetic disruption of *Frem1* results in reduced ano-genital distance, a phenotype also produced by genetic inactivation of *Gli1* transcription factors [4; 37]. Shh signaling is also active in skin development [1], where *Shh* expression in the epidermis initiates pathway activity in the epidermis and mesenchyme of the dermis. Shh pathway activity localizes to developing hair follicles, and its inhibition disrupts formation of new hair follicles [50]. *Frem1* expression appears more widespread in the early dermal mesenchyme but also progressively localizes to dermal papillae of hair follicles [46]. Our findings from facial morphogenesis argue that further investigation into whether Shh signaling regulates *Frem1* across other developmental contexts is warranted.

*Frem1* has been demonstrated to have multiple context-dependent roles during development. Apart from complexing with FREM2 and FRAS1 in dermis-epidermis adhesion, studies have identified a role for FREM1 in mesenchymal proliferation during mammalian diaphragm development and primary mesenchymal cell migration in sea urchin development [5; 23]. These biological processes are also required for cNCC development and facial morphogenesis. cNCCs are specified at the dorsal margins of the anterior neural folds, undergo epithelial-to-mesenchymal transition, migrate extensively, and then rapidly proliferate to form the majority of the connective tissue of the head and face. Disruptions in cNCC migration and proliferation contribute to facial malformations and dysmorphology [11; 54]. While not impacting cNCC migration, we found that addition of FREM1 to cNCCs resulted in a concentration-dependent increase in proliferation (Fig. 6). The concentration-dependence of this effect is notable given the regulation of *Frem1* by a secreted morphogen and the spatial gradient of *Frem1* expression during facial morphogenesis. Our finding that FREM1 promotes post-migrational cNCC proliferation provides a plausible cellular mechanism for the midfacial deficiency and asymmetry previously documented in *Frem1* mutant mice [52].

The evidence presented here suggests that the activity of *Frem1* in cNCC biology and facial morphogenesis is likely distinct from its well characterized role in epidermis development. We found that the cNCC mesenchymal domain of *Frem1* is expansive and extends substantially beyond the regions of *Frem2/Fras1* expression domains in the adjacent epithelial compartments. We also found that addition of FREM1 alone was sufficient to induce proliferation of cultured cNCCs. While *Fras1* expression is detectable at a low basal level of expression, *Frem2* is not detectable by qPCR in these cells (Fig. S2), suggesting that it is dispensable for the observed activity of FREM1 protein. The FREM/FRAS ternary complex binds with additional ECM glycoproteins, including ITGA8 and NPNT1 in several developmental contexts [22; 51]. Surprisingly, we found that while Shh pathway activity induced *Frem1*, addition of SHH ligand or activation of SMO in cNCCs resulted in downregulation of *Npnt* and *Itga8* (Fig. S3). Why Shh signaling suppresses expression of these genes while inducing *Frem1* is unclear but may be part of a mechanism that supports distinct FREM1 activity in cNCCs during facial morphogenesis.

The findings presented in this study reveal *Frem1* regulation as a novel mechanism by which Shh signaling controls facial morphogenesis. These results provide a developmental framework to understand *FREM1*-associated facial variation and the relationship of these outcomes to those resulting from genetic or environmental disruption of the Shh signaling pathway. These findings expand our understanding of the mechanisms that regulate facial morphogenesis and how their disruption contributes to human face shape variation and facial malformations.

## Materials and Methods

### Animal studies

This study was conducted in strict accordance with the recommendations in the Guide for the Care and Use of Laboratory Animals of the National Institutes of Health. The protocol was approved by the University of Wisconsin School of Veterinary Medicine Institutional Animal Care and Use Committee (Protocol No. 13–081.0). C57BL/6J mice were purchased from The Jackson Laboratory and housed under specific pathogen-free conditions in disposable, ventilated cages (Innovive, San Diego, CA). Rooms were maintained at 22 ± 2 °C and 30–70% humidity on a 12h our light, 12 hour dark cycle. Mice were fed 2920× Irradiated Harlan Teklad Global Soy Protein-Free Extruded Rodent Diet until day of plug, when dams received 2919 Irradiated Teklad Global 19% Protein Extruded Rodent Diet. One or two nulliparous female mice were placed with a single male for 1-2 hours and then examined for copulation plugs. The beginning of the mating period was designated as gestational day (GD)0, and pregnancy was confirmed by assessing weight gain between GD7 and GD10, as previously described [16].

Pregnant dams were administered 90 mg/kg/day cyclopamine (LC Laboratories, CAS #4449-51-8) or vehicle alone from GD8.25 to approximately GD9.375 by subcutaneous infusion exposure using ALZET 2001D micro-osmotic pumps (Cupertino, CA, United States) as previously described [10; 30]. Pregnant dams were euthanized by carbon dioxide inhalation followed by cervical dislocation for embryo collection at GD10.25 ± 1 h. Separation of mesenchyme and surface ectoderm of GD11 medial nasal processes and maxillary processes was accomplished as previously described [10; 28]. Tissues from an entire litter were pooled, and N=4 litters were used for RNA expression analyses. GD9.25 frontonasal prominence (FNP) tissue was microdissected and pooled as previously described [10]. N=6 litters for each treatment group were used for RNA expression analyses.

### In situ hybridization

Embryos at GD10, 10.25, 11, and 13 were dissected in PBS and fixed in 4% paraformaldehyde for 18 hours. Embryos then underwent graded dehydration (1:3, 1:1, 3:1 v/v) into 100% methanol and were stored at −20°C indefinitely for subsequent ISH analysis. Rehydrated embryos were embedded in 4% agarose gel and cut in 50 μm sections using a vibrating microtome. ISH was performed as previously described [14]. Sections were imaged using a MicroPublisher 5.0 camera connected to an Olympus SZX-10 stereomicroscope. Gene-specific ISH riboprobe primers were designed using IDT PrimerQuest and affixed with the T7 polymerase consensus sequence plus a 5-bp leader sequence to the reverse primer. Sequences are listed in Table S1.

### Cell culture

Immortalized O9-1 cranial neural crest cells were provided by Dr. Robert Maxson, Keck School of Medicine at the University of Southern California, and cultured as described by Ishii and colleagues [17]. O9-1 cell lines stably overexpressing GFP or a constitutively active form of Smoothened (SMO^M2^) were generated as previously described [10].

### Gene expression analysis

O9-1 cells were plated at 5 × 10^5^ cells/mL (0.4 mL per well in a 24-well plate) and were allowed to attach in complete O9-1 media for 24 hours. Media were replaced with DMEM containing 1% FBS and treatments of ± 0.4 μg/mL SHH ligand (R&D Systems) ± 100 nM vismodegib (LC Laboratories) or ± 50 nM Smoothened Agonist (LC Laboratories). Cells were harvested at 48 hours following treatment for RNA extraction. N=5 biological replicates were used in each treatment group. RNA was isolated from cells grown *in vitro* and from embryonic tissue using the Illustra RNAspin kit according to the manufacturer recommendations with on-column DNase digestion. cDNA was synthesized from 100-500 ng of total RNA using the GoScript reverse transcription reaction kits (Promega). Singleplex quantitative real-time polymerase chain reaction (qPCR) was performed using SSoFast EvaGreen Supermix (Bio-Rad) on a Bio-Rad CFX96 real-time PCR detection system (Bio-Rad Laboratories). qPCR primers were designed using PrimerQuest (IDT), and sequences are listed in Table S2. Target gene specificity was confirmed using National Center for Biotechnology Information Primer-Basic Local Alignment Search Tool (NCBI Primer-BLAST). *Gapdh* was used as the housekeeping gene, and analyses were conducted with the 2-ΔΔCt method.

### GLI binding site analyses

To identify GLI binding sites in the developing face, published FLAG ChIP-seq data from *Gli3*^*3XFlag*^ knock-in mice were accessed from GEO record GSE146961 [9; 32]. Raw sequencing reads were aligned to the mm10 genome using Bowtie2 and SAMtools [27] and filtered for a minimum mapping quality of 10 (-q 10). Fold-enrichment tracks were generated with BEDTools (v2.30.3) using the genomecov and bedGraphToBigWig functions [40]. Tracks were then visualized with the IGV genome browser (v2.4.14) [42] and edited with Adobe Illustrator (Adobe Inc., San Jose, CA).

### Migration assays

O9-1 cells were plated at 5 × 10^5^ cells/mL (0.4 mL per well in a 24-well plate) and allowed to attach in complete O9-1 media for 16 h. Immediately prior to beginning treatments, each well was scratched with a sterile 200 μL pipet tip vertically and washed twice with DPBS. Recombinant FREM1 protein (R&D Systems, Minneapolis, MN) dissolved in DPBS was diluted to final concentrations of 7.5 µg/mL, 2.5 µg/mL, and 0.83 µg/mL in DMEM containing 1% FBS, and 400 µL treatment or DPBS vehicle media were added to each well. Phase contrast images were taken with a MicroPublisher 5.0 camera (QImaging, Tucson, AZ) mounted on a Nikon Eclipse TS100 (Nikon Instruments Inc., Melville, NY) using a 4× objective every hour for 8 hours following the initiation of the treatment period. Migration rate was assessed using ImageJ. Experiments were performed in biological triplicate.

### Proliferation assays

O9-1 cells were plated at 5 × 10^4^ cells/mL (0.4 mL per well in a 24-well plate) in Matrigel-coated wells and allowed to attach in complete O9-1 media for 16 h. Recombinant FREM1 protein (R&D Systems, Minneapolis, MN) dissolved in DPBS was diluted to final concentrations of 7.5 µg/mL, 2.5 µg/mL, and 0.83 µg/mL in DMEM containing 1% FBS, and 400 µL treatment or DPBS vehicle media were added to each well. At 22 h post-treatment, half of the media were removed from each well and replaced with a 2× solution of EdU (Invitrogen, Waltham, MA), per the manufacturer’s recommendations. Cells were incubated in the EdU solution for 2 hours prior to fixation in 4% paraformaldehyde for 15 min. EdU and Hoechst staining were performed according to the manufacturer’s instructions with volumes adjusted for use in a 24-well plate. Fluorescent images were captured using the Keyence BZ-X700 fluorescent microscope (Keyence, Itasca, IL), and Hoechst+ and EdU+ cells were counted in 4 non-adjacent fields per well using Keyence BZ-X Image Analysis software. Experiments were performed in technical duplicate for a total of N=3 biological replicates.

### Statistics

One-way analysis of variance (ANOVA) with Tukey’s post hoc test or two-tailed t-tests, where appropriate, were used to determine whether gene expression was changed in cultured cNCCs and in microdissected FNP tissue. ANOVA with Dunnett’s post hoc test for multiple comparisons was used for analyses of cNCC migration and proliferation assays. GraphPad Prism 6 was used for all statistical analyses. An alpha value of 0.05 was maintained for determination of significance.

## Acknowledgments

Research reported in this publication was supported by the National Institute of Dental and Craniofacial Research and the National Institutes of Health under award numbers R00DE022010, R03DE027162, and T32ES007015. The content is solely the responsibility of the authors and does not necessarily represent the official views of the National Institutes of Health. Support was also provided by a University of Wisconsin Hilldale Undergraduate Research Award. The authors thank Drs. Ian Smyth, Daryl Scott, and Kiyotoshi Sekiguchi for their generous willingness to share reagents and Dr. Suzanne Ponik for helpful discussion.

## Notes

### Competing Interest Statement

The authors have declared no competing interest.

